# Strain-level variation influences *Methanothermococcus* fitness in marine hydrothermal systems

**DOI:** 10.1101/2021.05.29.446292

**Authors:** Michael Hoffert, Rika Anderson, Julie Reveillaud, Leslie Murphy, Ramunas Stepanauskas, Julie A. Huber

**Author notes:** These authors contributed equally to this paper (Michael Hoffert, Rika Anderson).

## Abstract

Hydrogenotrophic methanogens are ubiquitous chemoautotrophic archaea inhabiting globally distributed deep-sea hydrothermal vent ecosystems and associated subseafloor niches within the rocky subseafloor, yet little is known about how they adapt and diversify in these habitats. To determine genomic variation and selection pressure within methanogenic populations at vents, we examined five *Methanothermococcus* single cell amplified genomes (SAGs) in conjunction with 15 metagenomes and 10 metatranscriptomes from venting fluids at two geochemically distinct hydrothermal vent fields on the Mid-Cayman Rise in the Caribbean Sea. We observed that some *Methanothermococcus* strains and their transcripts were more abundant than others in individual vent sites, and among the genes differentiating the *Methanothermococcus* SAGs were those related to nitrogen fixation and the CRISPR/Cas immune system. SAGs possessing these genes were less abundant than those missing that genomic region. Overall, patterns in nucleotide variation indicated that the population dynamics of *Methanothermococcus* were not governed by clonal expansions or selective sweeps, at least in the habitats and sampling times included in this study. Together, our results show that although specific lineages of *Methanothermococcus* co-exist in these habitats, some outcompete others, and possession of accessory metabolic functions does not necessarily provide a fitness advantage in these habitats in all conditions. This work highlights the power of combining single-cell, metagenomic, and metatranscriptomic datasets to determine how evolution shapes microbial abundance and diversity in hydrothermal vent ecosystems.

## Introduction

Hydrothermal vents host diverse ecosystems sustained by physical and geochemical gradients that are created by the mixing of high-temperature hydrothermal fluid with cold seawater. Several investigations have revealed the intimate relationships between geochemistry, microbial community structure, and metabolic processes in these systems (Huber et al., 2007; Flores et al., 2012; Akerman et al., 2013; Perner et al., 2013; Reveillaud et al., 2016; Meier et al., 2017; Fortunato et al., 2018). However, little is known about how natural selection molds the genomes of metabolically and taxonomically diverse microbial species inhabiting the rocky subseafloor near hydrothermal vents, especially with respect to genomic variation and plasticity at the population level.

Microbial genomes are highly variable, facilitated by extensive gain, loss, and horizontal transfer of genes between microbes. Previous studies have shown that genome plasticity within species can allow for adaptation to diverse environments (Medini et al., 2005; Tettelin et al., 2005), including in hydrothermal vent habitats (Marteinsson et al., 1999; White et al., 2008; Brazelton et al., 2011; Meyer and Huber, 2014; Moulana et al., 2020). The accessory genome in vent microbial species has been found to include genes related to nitrogen metabolism (Meyer and Huber, 2014; Galambos et al., 2019) and nutrient uptake (Moulana et al., 2020), as well as glycosyltransferases, S-layer proteins, and other membrane-related genes (Anderson et al., 2017). Some studies suggest that the adaptation of particular microbial strains to specific environmental conditions may enable clonal expansions of these strains (Bendall et al., 2016; Shapiro, 2016). Investigation of genomic variation in vents identified patterns of genomic variation indicative of such clonal sweeps (Anderson et al., 2017).

In hydrothermal vent systems, methanogenic archaea are abundant in the subseafloor, venting fluids and the interiors of active chimneys in both basalt- and serpentinite-hosted systems (Schrenk et al., 2003; Takai et al., 2004; Brazelton et al., 2006; Ver Eecke et al., 2013). These archaea consume hydrogen and carbon dioxide, formate, or acetate to produce methane and can grow across a range of temperatures (Takai et al., 2008; Thauer et al., 2008; Stewart et al., 2019). Methanogenic archaea can vary their genomic repertoire in response to carbon sources like formate (Hansen et al., 2011), and single-species methanogenic biofilms from carbonate chimneys at vents show evidence of physiological differentiation (Brazelton et al., 2011).

At the Mid-Cayman Rise, an ultra-slow spreading ridge located in the Caribbean Sea, methanogens from the genus *Methanothermococcus* comprised the most abundant lineage of archaea observed in venting fluids (Reveillaud et al., 2016). The Mid-Cayman Rise hosts two geologically and spatially distinct vent fields: Piccard vent field, an ultra-deep (4960m) hydrothermal system situated in mafic rock, and Von Damm vent field, a shallower (2350m) ultramafic site on a nearby massif (German et al., 2010). The Von Damm hydrothermal field features fluids enriched with H_2_ and CH_4_ produced during serpentinization of the ultramafic bedrock in which it is situated (German et al., 2010; Hodgkinson et al., 2015; McDermott et al., 2015, 2020). In contrast, the mafic Piccard hydrothermal field is characterized by fluids with elevated concentrations of sulfide and hydrogen at lower pH and higher temperature (German et al., 2010). Previous work characterizing microbial communities in venting fluids from both vent fields at Mid-Cayman Rise demonstrated that although microbial community structure varies both within and between vent sites, the microbial communities exhibit functional redundancy across sites (Reveillaud et al., 2016). Later work revealed differences in metabolic gene expression across vent sites at the Mid-Cayman Rise, with genes related to methanogenesis more transcribed at Von Damm (Galambos et al., 2019). Patterns of fine-scale genomic variation indicate population-specific differences in selection pressure between Von Damm and Piccard (Anderson et al., 2017). More globally, the ubiquitous sulfur-oxidizing Epsilonbacteraeota genus *Sulfurovum* was shown to have an extensive pangenome, with distinct geographic patterns in gene content between *Sulfurovum* strains in the Mid-Cayman Rise and Axial Seamount in the Pacific Ocean (Moulana et al., 2020). Similar strain-level variation could occur in methanogenic archaea at the Mid-Cayman Rise. 16S rRNA gene sequencing found that different OTUs of *Methanothermococcus* display varying abundance across samples, indicating niche partitioning within vent sites and between vent fields (Reveillaud et al., 2016). However, niche partitioning and genomic variation have not been examined at the genome scale for thermophilic methanogens, nor is it known which genes have the strongest influence on methanogen fitness.

To gain insight into genomic variation among methanogens inhabiting hydrothermal systems, we examined five single-cell amplified genomes (SAGs) from the genus *Methanothermococcus* in conjunction with 15 metagenomes and 10 metatranscriptomes from the Von Damm and Piccard vent fields at the Mid-Cayman Rise. We examine how the differential abundance of *Methanothermococcus* lineages and their transcripts correlate with the presence and absence of particular genes. Investigation of single nucleotide variation across SAGs and across gene types provides further insights into the way differential gene content can influence fitness of this globally distributed microbial group in hydrothermal vent environments.

## Materials and Methods

### Sample collection

We collected diffuse flow hydrothermal fluid samples during cruises aboard the R/V *Atlantis* in January 2012 (FS841-FS856) and the R/V *Falkor* in June 2013 (FS866-FS881) using the ROV *Jason* and HROV *Nereus*. Further information about sample collection can be found in Reveillaud et al. (2016) and Galambos et al. (2019). Samples for single-cell genomics analysis were collected by preserving 1 mL of hydrothermal vent fluid with 100µl glyTE solution (4 parts 11x TE to 5 parts glycerol (Stepanauskas et al., 2017) followed by incubation for 5 min. at room temperature and storage at -80°C. For metagenomics and metatranscriptomics, we pumped ∼3-6 liters of fluid through 0.22 µm Sterivex filters (Millipore) on the seafloor, flooded the filters with RNALater (Ambion) shipboard, sealed the filters with Luer Caps and stored them in sterile Falcon tubes for 18 hours at 4°C, then froze them at -80°C.

### Single-cell genomics

Single cell genomics analyses were performed on one diffuse fluid sample from Ginger Castle site at the Von Damm vent field (sample FS848; see Table 1). The field sample was thawed, stained with SYTO-9 nucleic acids stain (Thermo Fisher Scientific) and pre-filtered through a 40 µm mesh-size cell strainer (Becton Dickinson); the individual cells were separated using fluorescence-activated cell sorting (FACS), lysed using a combination of freeze-thaw and alkaline treatment, their genomic DNA was amplified using multiple displacement amplification in a cleanroom, and the resulting single amplified genomes (SAGs) were screened for the 16SrRNA genes as previously described (Stepanauskas et al., 2017). This work was performed at the Bigelow Laboratory Single Cell Genomics Center (https://scgc.bigelow.org; SCGC).

**Table 1.** Summary of sequencing and assembly statistics for five *Methanothermococcus* single cell genomes isolated from diffuse fluids at Ginger Castle in Von Damm hydrothermal field.

| Name | Completeness (%) | Number of scaffolds >1 kbp | Assembly size (bp) | GC content (%) | Number of ORFs | N50 |
| --- | --- | --- | --- | --- | --- | --- |
| SCGC AD-155-C09 | 84.27 | 125 | 1,740,890 | 29.10 | 1474 | 25,888 |
| SCGC AD-155-E23 | 80.5 | 51 | 1,043,924 | 42.97 | 1221 | 56,319 |
| SCGC AD-155-K20 | 16.01 | 25 | 188,232 | 30.59 | 205 | 8,893 |
| SCGC AD-155-M21 | 67.77 | 107 | 932,247 | 30.37 | 937 | 17,756 |
| SCGC AD-155-N22 | 55.15 | 99 | 856,655 | 35.21 | 866 | 14,681 |

Based on 16S rRNA gene analysis, 5 *Methanothermococcus* SAGs were chosen for sequencing. Libraries were prepared using the Ovation Ultralow Library DR multiplex system (Nugen) and were sheared at 250bp using a Covaris S-series sonicator. All SAGs were sequenced with NextSeq 500 (Illumina) at the W.M. Keck Facility in the Josephine Bay Paul Center at the Marine Biological Library. Each SAG was sequenced on a different plate to reduce the possibility of cross-contamination through “index-switching” or “sample bleeding” (Mitra et al., 2015; Sinha et al., 2017).

Reads were trimmed and merged using the illumina-utils software package (Eren et al., 2013). To maximize contig length while minimizing the risk of chimeras, reads were assembled using 4 different software packages: IDBA-UD v.1.1.2 (Peng et al., 2012), SPAdes 3.10.0 (Bankevich et al., 2012), CLC Genomics Workbench v. 7, and A5 (Coil et al., 2015). Contigs from all four assemblers were combined, compared for redundancy, and integrated into a new assembly using the Contig Integrator for Sequence Assembly of bacterial genomes (CISA) (Lin and Liao, 2013).

### Pangenomic analysis of single cell genomes

A phylogenetic tree of all SAGs was constructed using RaxML (Stamatakis, 2014) with 100 bootstraps, using the PROTGAMMAAUTO flag to allow the program to determine the best protein model of evolution. The alignment was created from universal single copy marker genes that were identified and aligned using Phylosift (Darling et al., 2014). The reference sequences were obtained from NCBI, with the following accession numbers: *Methanococcus maripaludis* C7 [NCBI:txid426368] (designated as the outgroup), *Methanothermococcus thermolithotrophicus* DSM 2095 [NCBI:txid523845], and *Methanothermococcus okinawensis* IH1 uid51535 [NCBI:txid647113].

Open reading frames were identified and annotated for all SAGs using the JGI IMG/ER pipeline (Markowitz et al., 2012). Visualizations of ORFs on contigs and cross-links between ORFs were created using the Biopython collection (Cock et al., 2009) with the GenomeDiagram module. Clustering of ORFs was conducted using the Integrated Toolkit for the Exploration of microbial Pangenomes (ITEP) (Benedict et al., 2014). We clustered the ORFs using an inflation value of 2 and a maxbit score of 0.3. ITEP was also used to compare ORF presence/absence across SAGs. KEGG Decoder (Graham et al., 2018) was used to determine pathway completeness for selected metabolic pathways, and the heatmap was visualized with the Python Seaborn library (Waskom et al., 2014). PAML v4.9 (Yang, 2007) was used to calculate dN/dS values for each cluster with greater than one gene assignment. We compiled the results of PAML analysis and the presence/absence table using a custom Python (v.2.7.5) script available on GitHub at https://github.com/carleton-spacehogs/SAGs.

Reads from each SAG were mapped to the contigs of each of the other SAGs using bowtie2 v. 2.2.9 (Langmead and Salzberg, 2012) using default settings. Coverage was calculated using samtools v1.3.1 (Li et al., 2009), and bedtools v2.26.0 (Quinlan, 2014). Average nucleotide identity was calculated using the software package pyani v.0.2.7 (Pritchard et al., 2016).

### Microbial community DNA and RNA preparation and sequencing

To extract DNA from Sterivex filters for sequencing, we followed methods described in Reveillaud et al. (2016) and Galambos et al. (2019). We extracted total genomic DNA from half of the Sterivex filter, and we extracted RNA from the other half of the Sterivex filter using the mirVana miRNA isolation kit (Ambion), adding a bead-beating step using beads from the RNA PowerSoil kit (MoBio) as described in Fortunato and Huber (2016). RNA was treated with DNAse using the Turbo-DNase kit (Ambion), and then purified and concentrated using the RNAeasy MinElute kit (Qiagen). We prepared metatranscriptomic libraries using the Ovation Complete Prokaryotic RNA-Seq DR multiplex system (Nugen) following the manufacturer’s instructions. We prepared the metagenomic libraries as described in Reveillaud *et al*. (2016). All libraries were sheared at 175bp using a Covaris S-series sonicator, yielding paired-end reads with a 30bp overlap. We sequenced all libraries on an Illumina Hi Seq 1000 at the W.M. Keck Facility in the Josephine Bay Paul Center at the Marine Biological Laboratory.

We merged and filtered metagenomic and metatranscriptomic reads using the illumina-utils package (Eren et al., 2013) using iu-merge-pairs with the –enforce-Q30-check flag, followed by iu-filter-merged-reads with a maximum mismatch of 2 in the overlap region. This resulted in reads averaging approximately 170bp in length. We removed rRNA *in silico* from metatranscriptomes by mapping all merged, filtered reads to the Silva LSU and SSU Parc databases (release 111) (Pruesse et al., 2007; Quast et al., 2013) using bowtie2 v.2.2.9 with local alignment (Langmead and Salzberg, 2012) and removing all reads that mapped.

### Metagenome and metatranscriptome mapping

We mapped the metagenomic and metatranscriptomic reads (Supplementary Table 1) of each sample to the contigs of each SAG (Table 1, Supplementary Table) using bowtie2 v2.2.9 (Langmead and Salzberg, 2012) with default settings. We used anvi’o v6.1 (Eren et al., 2015) to identify, quantify and characterize read coverage, single nucleotide variants (SNVs) and single amino acid variants (SAAVs) for all of the metagenomic reads mapping to each of the SAGs. SNVs and SAAVs were only counted if they had a minimum coverage of 10, and SAAVs were only counted if the read matching the variant mapped to the entire codon context. Anvi’o was also used to calculate average mapping coverages of metagenomes and metatranscriptomes for each of the SAGs. A custom Python database of ORFs, contigs, SNVs, SAAVs, and clusters was compiled for cross-referencing annotation, variability, and homologous clustering data. From this database, SNV densities were calculated based on the length of annotated ORFs, and underlying distributions of SNV density for each sample were estimated using the Python Seaborn library (Waskom et al., 2014). Plots for SNV density vs. n2n1 (or the ratio of the second most frequent nucleotide to the consensus nucleotide), coverage, bubble plots, and average abundance were generated using lookups in this database on a sample- or genome-specific basis. SNV densities were matched to genes annotated by the JGI-IMG annotation pipeline (Markowitz *et al*., 2012) using in-house Python scripts, available on GitHub at https://github.com/carleton-spacehogs/SAGs. Box-and-whisker plots depicting metatranscriptome coverage of specific genes were created using the Python Seaborn library (Waskom et al., 2014).

### CRISPR and prophage identification

CRISPR loci were identified in SAG contigs using the CRISPR Recognition Tool (CRT) (Bland et al., 2007). Prophage sequences were identified in SAG contigs using VirSorter (Roux et al., 2015) on the CyVerse platform with default settings.

### Data availability

This Whole Genome Shotgun project has been deposited at DDBJ/ENA/BenBank under accessions All assembled SAG contigs were deposited at NCBI under accessions JAHEQB000000000 (C09), JAHEQA000000000 (E23), JAHEPZ000000000 (K20), JAHEPY000000000 (M21), and JAHEPX000000000 (N22). The versions described in this paper are versions JAHEQB010000000 (C09), JAHEQA010000000 (E23), JAHEPZ010000000 (K20), JAHEPY010000000 (M21), and JAHEPX010000000 (N22). Reads for each SAG were deposited at ENA under accession numbers ERS6485300, ERS6485304, ERS6485303, ERS6485302, and ERS648530. Metagenomic and metatranscriptomic reads are deposited under study accession code PRJEB15541 in the EMBL-EBI European Nucleotide Archive (ENA) database. All code used for this project has been deposited to GitHub at https://github.com/carleton-spacehogs/SAGs.

## Results

### Characterization of single-cell amplified genomes (SAGs)

Single-cell amplified genomes (SAGs) were generated from ∼47°C fluids from the Ginger Castle vent site at Von Damm in 2012 (Reveillaud et al., 2016) (Supplementary Table 1). A summary of the sequencing and assembly of the SAGs is presented in Table 1. The SAGs are labeled with their full names in Table 1; for the remainder of the manuscript, we will refer to them simply as C09, E23, K20, M21, and N22. The N50 of the assemblies ranged from 56,319 - 8,893 bp, and assembly completeness based on the presence of single-copy universal markers ranged from 84% to 16%. A phylogenetic tree constructed from universal single-copy marker genes revealed two distinct clades, one containing C09, M21, and K20, and the other containing N22 and E23 (Figure 1). We mapped the reads of each SAG to the contigs of every other SAG to identify shared genomic regions, and the degree of shared genomic content generally matched the phylogenetic relationships as determined by single copy universal genes (Supplementary Table 2). The five SAGs showed low mapping coverage to one another in general (∼9% reads mapping on average), but we observed relatively higher average coverage in SAG-to-SAG mappings among C09, M21, and K20 (∼30% reads mapping on average). Calculation of average nucleotide identity (ANI) revealed that C09, M21 and K20 were most similar to each other (approximately 94% ANI on average), whereas E23 and N22 were more distantly related to all the others (approximately 84% ANI to all the other SAGs on average). This is consistent with the phylogenetic relationships illustrated by single-copy universal genes (Figure 1).

**Figure 1.**
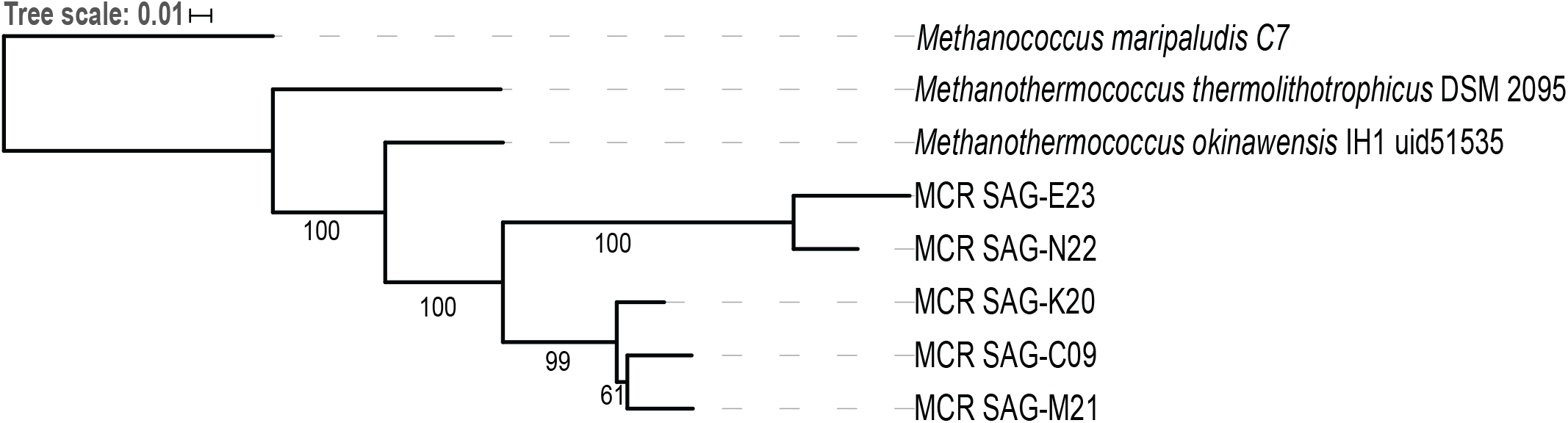
Phylogenetic tree of five *Methanothermococcus* SAGs from the Mid-Cayman Rise (MCR). Single-copy universal marker genes were identified and aligned with PhyloSift. The tree was built with RAxML using 100 bootstraps. Bootstrap values higher than 50 are indicated on the tree. Each of the SAGs sequenced for this study was labeled as “MCR SAG” in the tree for clarity.

### *Gene and transcript abundance of* Methanothermococcus *lineages across diffuse sites*

To determine whether the lineages represented by these *Methanothermococcus* SAGs varied in gene and transcript abundance among vent sites, we mapped reads from 15 metagenomes and 10 metatranscriptomes from different sites in the Piccard and Von Damm vent fields to each SAG (Figure 2). The SAGs had highest coverage in Von Damm vent sites, including Ginger Castle (FS848), Near Main Orifice (FS866), Shrimp Buttery (FS877), and Old Man Tree (FS881). With the exception of Ginger Castle (48°C), which is the site where the SAGs where sampled from, these vents were characterized by high-temperature fluids (114°C-131°C, Supplementary Table 1). We observed substantial variation in the relative abundance of the lineages represented by these SAGs between sample sites, with E23 and N22 significantly more abundant compared to the other SAG lineages at two vent sites in particular, Old Man Tree and Near Main Orifice. In these sites, E23 had the highest coverage of all the SAGs (Figure 2A).

**Figure 2.**
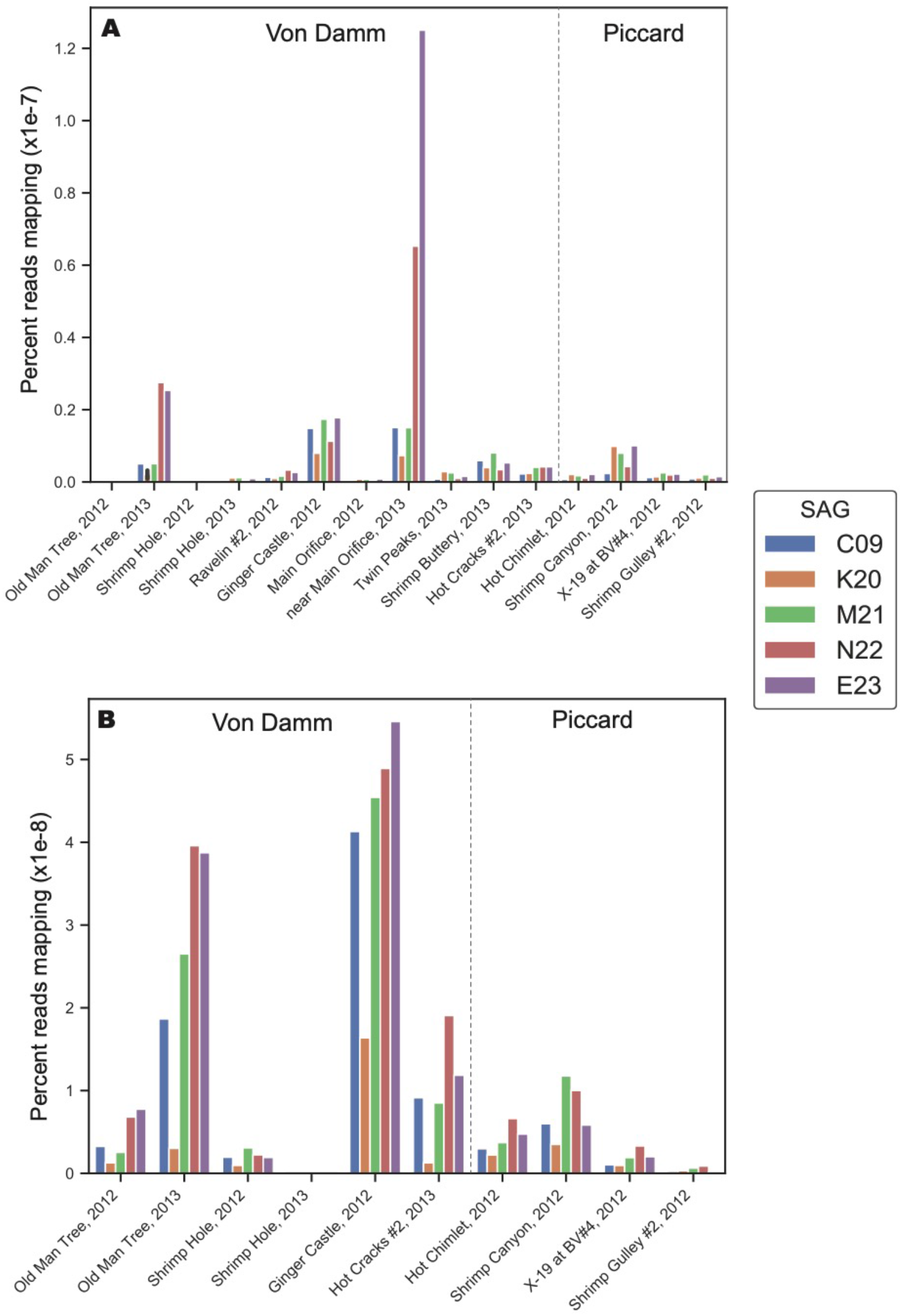
Normalized percent mapping of A) metagenomic reads and B) metatranscriptomic reads for each SAG assembly. The total number of mapped reads to each SAG was divided by the number of metagenomic or metatranscriptomic reads and by total SAG length. Supplementary Table 1 lists all samples used for mapping.

The differences in metagenomic coverage among the SAGs were reflected in differences in transcript abundance, which is an indicator of gene expression. At the whole-genome level, the *Methanothermococcus* lineages represented by these SAGs had the highest transcript abundance at Ginger Castle, the site where the SAGs were sampled from (Figure 2B). The lineages represented by SAGs E23 and N22, which share a clade distinct from the other *Methanothermococcus* SAGs (Figure 1), consistently showed the highest transcript abundance, especially at Old Man Tree and Hot Cracks #2 in the Von Damm vent field. This is roughly consistent with their coverage based on metagenomic mapping (Figure 2A).

### *Analysis of gene presence and absence across* Methanothermococcus *SAGs*

To determine whether variation in gene content drove the observed differences in gene and transcript abundance, we examined variable gene content across each of the SAGs. To search for shared genes across all SAGs based on sequence similarity, we clustered all ORFs based on all-v-all BLAST hit scores and compared the presence and absence of gene clusters across each SAG. It is important to note that these SAGs varied in completeness, and thus, absence of evidence for specific genes does not constitute evidence of absence. Therefore, very few genes were considered “core” (that is, found in all five SAGs). Rather than focusing on individual gene presence/absence trends, we instead searched for consistent patterns in gene presence/absence that shared similar functions across the five SAGs. We focused on groups of functional genes whose presence/absence patterns match the phylogenetic relationships shown in Figure 1.

Homologous clustering of the open reading frames (ORFs) from all five SAGs produced 1,838 clusters of homologous ORFs. ORFs with identical functional annotations and identical presence/absence patterns were tabulated and summarized (Supplementary Data File 1). Most variable genes were annotated as hypothetical proteins. The SAGs contained several ORFs that encoded functional genes related to the metabolism of hydrogen, sulfur, and nitrogen (Supplementary Table 3), many of which showed consistent presence/absence patterns. All five SAGs contained genes for iron transport proteins, energy-converting hydrogenases, fumarate reductase, and methyl coenzyme M. Analysis of metabolic pathway completeness revealed that all five SAGs encoded complete or nearly complete pathways for methanogenesis via CO_2_ and acetate, but lacked the genes required for methanogenesis via dimethylsulfide, methanethiol, or methylamines (Figure 3). Surprisingly, we did not observe formate dehydrogenase genes in any of the *Methanothermococcus* SAGs. N22 uniquely encoded genes related to chemotaxis that were missing in the other SAGs (Figure 3). K20 was missing many metabolic pathways, and had largely incomplete pathways, most likely due to the incompleteness of the SAG assembly (16%).

**Figure 3.**
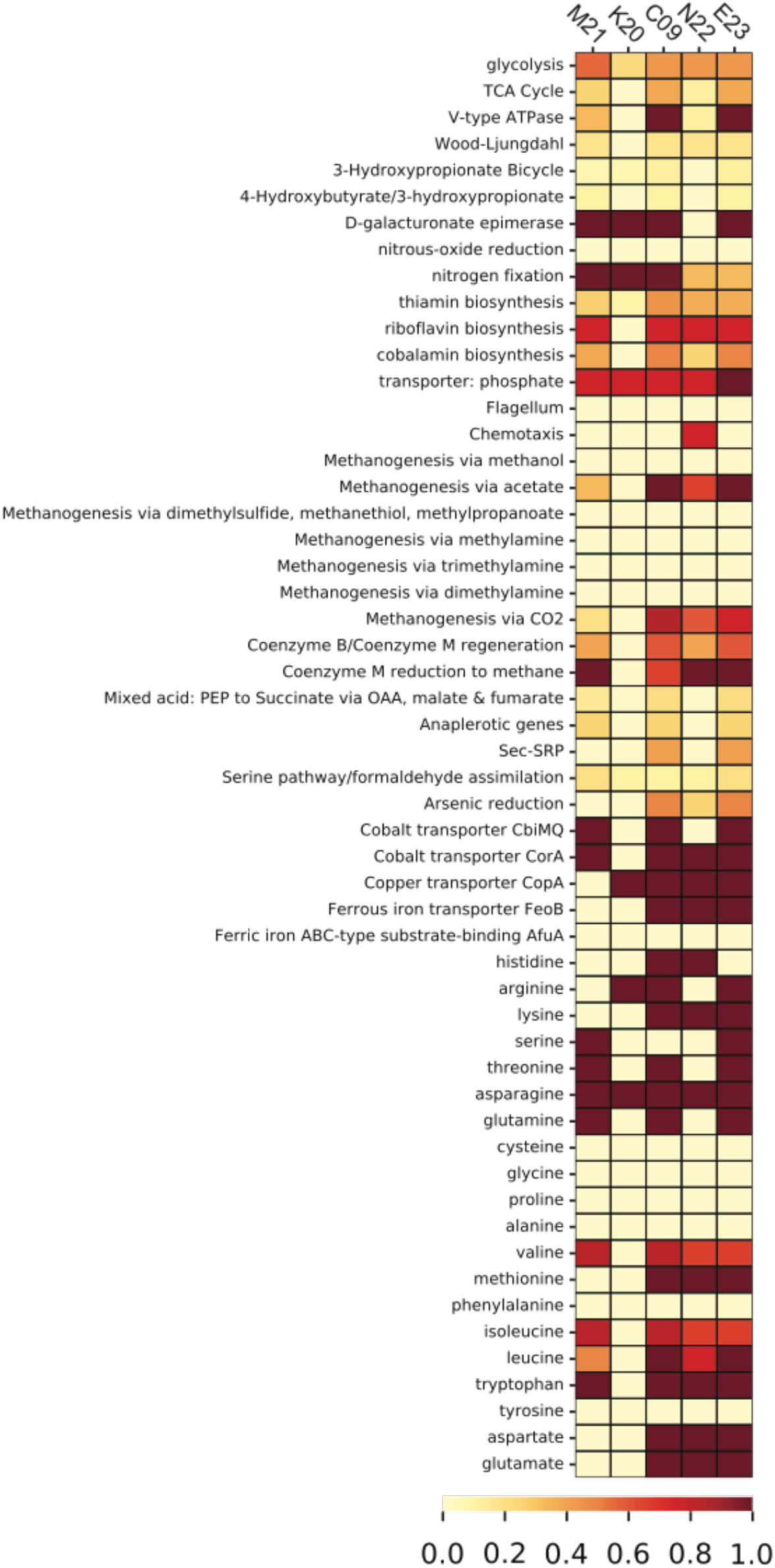
Heatmap depicting pathway completeness of selected metabolic pathways in each of the five *Methanothermococcus* SAGs. Darker values indicate more complete pathways.

We identified several genes whose presence/absence patterns matched the SAG phylogenetic relationships. SAGs C09 and M21 shared a large set of gene clusters, which matches their placement on the phylogenetic tree (Figure 1), including a cluster of CRISPR-associated proteins (Supplementary Table 3). The abundance of genes shared within this clade was in direct contrast to E23 and N22, a clade distinguished primarily by the absence of unique gene clusters relative to the C09/M21/K20 clade (Supplementary Table 3, Supplementary Data File 1). We identified a genomic region encoding several genes related to nitrogen metabolism, including *nif*, that was present in genomes C09/K20/M21 but not in E23 or N22, consistent with phylogenetic clustering according to single-copy universal genes (Figure 4A, Supplementary Table 3). Assessment of metabolic pathway completeness in each SAG confirmed that E23 and N22 encoded incomplete nitrogen fixation pathways, while C09/K20/M21 all encoded complete nitrogen fixation pathways (Figure 3). Synteny analyses revealed that the order of genes was consistent across SAGs C09/M21/K20 (Figure 4A). The conserved genes on this island include the alpha and beta chains of a nitrogenase molybdenum-iron protein, a molybdate transport system, and the nitrogen regulatory protein PII. We identified this genomic island on three separate contigs in K20, and although BLAST comparison indicated that the island was not identical between contigs, we cannot eliminate the possibility that the placement of these sequences on three separate contigs resulted from a mis-assembly and actually represents a single region on the genome. This genomic region is surrounded by genes related to glycosyltransferases, methyltransferases, asparagine synthase and tRNA synthetase, and although similar sequences of ORFs were identified in SAG E23, the conserved set of nitrogenase genes identified in C09/M21/K20 were not found on this contig (Figure 4A), further suggesting that the island was missing from the E23 genome and not simply missing from the SAG sequence due to incomplete sequencing or assembly.

**Figure 4.**
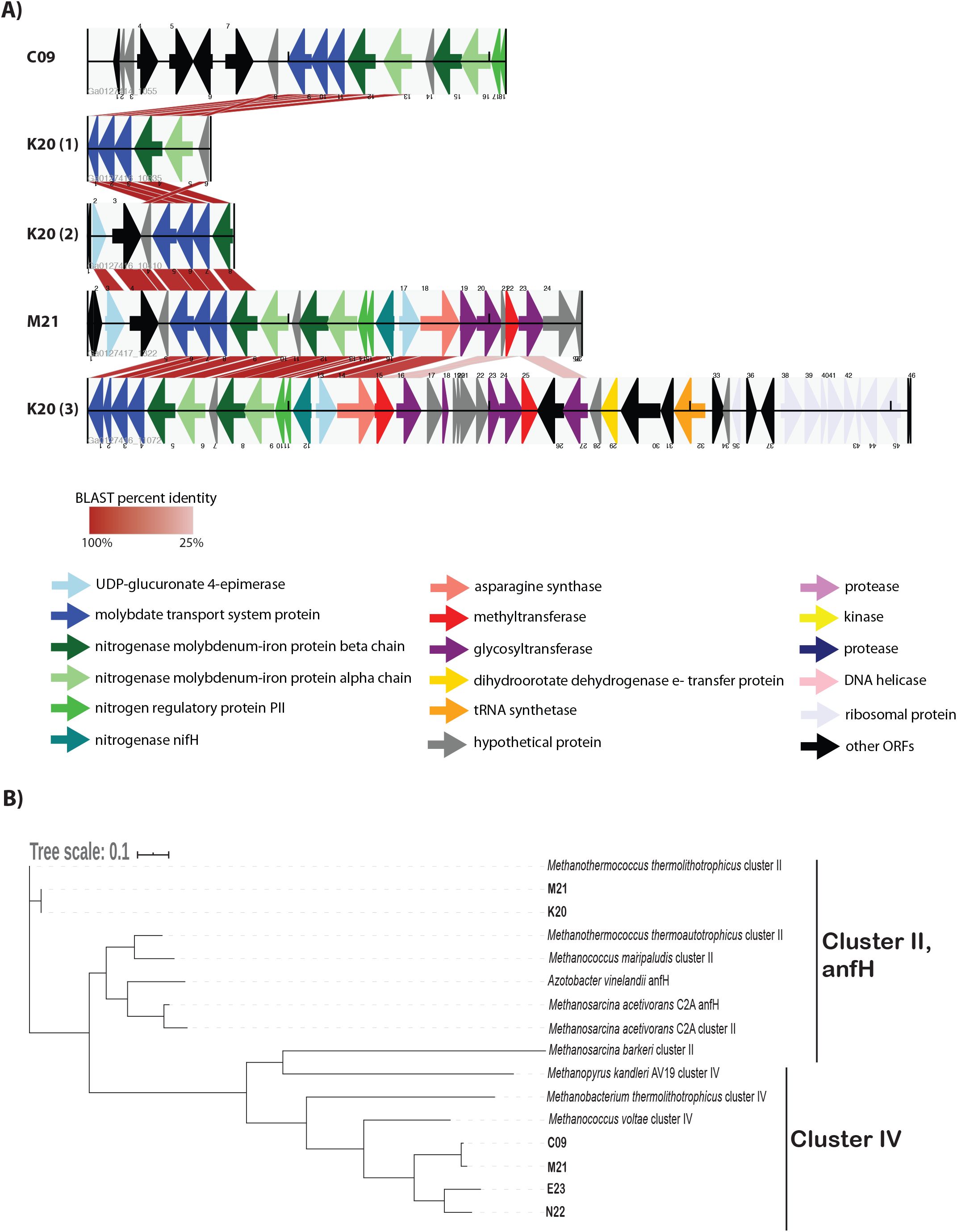
A) Alignment of open reading frames associated with nitrogenase genes encoded in *Methanothermococcus* SAGs. The open reading frames (ORFs) are represented by arrows indicating the direction of transcription. ORFs are color-coded according to their predicted function as determined by the JGI IMG annotation pipeline. Cross-links indicate the best match to the ORFs on the corresponding contig as determined by BLAST analysis; shading of the cross-link indicates the BLAST percent identity, with darker red colors corresponding to higher percent identity. B) Maximum likelihood tree depicting phylogenetic relationships among nifH genes identified in *Methanothermococcus* SAGs. SAG gene numbers are included in the SAG leaf name, and GenBank accession numbers for reference sequences are included in the Methods. The tree depicts sequences from cluster II and cluster IV nitrogenases. Note that the cluster II nitrogenases included in the tree appear as teal arrows in the ORF diagram. The tree was constructed with RAxML with 100 bootstraps.

While all of the SAGs encoded nifH-like genes, most of these nitrogenases fell into a clade containing cluster IV nitrogenases on a phylogenetic tree, which include divergent nitrogenases from archaea that cannot fix N_2_ (Chien and Zinder, 1994) (Figure 4B). Only the *Methanothermococcus* SAGs K20 and M21 encoded nifH genes from cluster II (Figure 4B), a clade which is known to include functional nitrogenases. These cluster II nifH genes were part of the genomic region that was conserved among C09, K20 and M21(Figure 4A).

There is evidence that nitrogenase genes were expressed in the *Methanothermococcus* strains encoding them. We observed elevated transcript abundance for the nitrogenase genes in SAGs C09, M21, and K20 at Ginger Castle in Von Damm (where these SAGs were isolated), though the transcript abundance for the nitrogenases was not as high as for genes related to methanogenesis (Supplementary Figure 1). Finally, although N22 was distinguished from other SAGs by the presence of chemotaxis-related genes on its genome, we observed zero transcripts mapping to these chemotaxis genes at Ginger Castle in Von Damm.

### CRISPR loci and prophage sequences

To examine viral-driven genomic differentiation among the SAGs, we searched for CRISPR loci and integrated prophage. Several groups of CRISPR-related proteins as well as CRISPR loci were identified in C09, M21, and K20. C09 and M21 each contained six CRISPR loci, with a total of 237 and 109 spacers, respectively, and K20 contained one CRISPR locus with just two spacers (Table 2, Supplementary Figure 2). No CRISPR genes or CRISPR loci were observed in E23 or N22. We observed substantial variation in spacer content among the CRISPR loci. The vast majority of CRISPR loci in C09, M21 and K20 were unique, with only 18 spacers matching between C09 and M21 (using an e-value cutoff of 0.001 to constitute a “match”, Supplementary Figure 2). These spacer matches did not occur in the same order, and occasionally matched other spacers within the same SAG. Moreover, many of these spacers matched CRISPR spacers from the cultured isolate *Methanothermococcus okinawensis* (Supplementary Figure 2). CRISPR spacers in SAGs C09 and M21 had matches to contigs in several of the metagenomes, with the most matches to sites in Von Damm, including Ginger Castle (FS848), near Main Orifice (FS866), Shrimp Buttery (FS877), Hot Cracks #2 (FS879), and Old Man Tree (FS881) (Table 2). Transcripts related to CRISPRs were detected at relatively low levels in SAGs C09 and M21 at Ginger Castle vent, where they were originally isolated (Supplementary Figure 1).

**Table 2.** Prophage and CRISPR loci in *Methanothermococcus* SAGs from the Mid-Cayman Rise (MCR). Phage clusters listed are as defined by the VirSorter database (Roux et al., 2015). BLAST matches correspond to a maximum e-value cutoff of 1e-05, unless otherwise stated. *M. okinawensis* phage were identified using VirSorter.

| CRISPR Loci |  |  |
| --- | --- | --- |
|  | Number of CRISPR Loci | Number of matches to MCR metagenome contigs |
| C09 | 6 | FS848 (10); FS854 (1); FS866 (26); FS877 (1); FS879 (4); FS881 (8) |
| E23 | 0 | -- |
| K20 | 1 | 0 |
| M21 | 6 | FS848 (1); FS866 (4); FS879 (1) |
| N22 | 0 | -- |
| Prophage |  |  |
|  | Prophage genes identified | Number of matches to *Methanothermococcus okinawensis* phage |
| C09 | Category 2 putative phage on 2 contigs. 27 genes total.<br>• 20 hypothetical proteins<br>• Terminase-like_family_protein<br>• phage putative head morphogenesis protein, SPP1 gp7 family<br>• phage protein, HK97 gp10 family<br>• phage tail tape measure protein<br>• phage major capsid protein E<br>• Bacteriophage lambda head decoration protein D | 22 |
| E23 | None | -- |
| K20 | None | -- |
| M21 | None | -- |
| N22 | None | -- |

In addition to the CRISPR sequences, several putative prophage sequences were identified in *Methanothermococcus* SAG C09. These genes contained phage-like proteins related to phage tail and phage head proteins, many of which match several prophage genes on the previously-sequenced *Methanothermococcus okinawensis* (e-value cutoff 1E-05) (Table 2). Genes in the integrated prophage region did not appear to be expressed at the time of sampling, as metatranscriptomic mapping to C09 indicated that the transcripts for prophage genes identified in C09 were present at low levels (with only ∼36 reads mapping to the prophage region 1 and ∼19 mapping to prophage region 2) at Ginger Castle in Von Damm (the site from which the SAGs were originally sampled).

### Selective signatures between SAGs

The dN/dS ratio, or ω, is generally used as an indicator to quantify the strength of selection by comparing the rate of substitution of synonymous sites (dS), which are ostensibly neutral, to the rate of nonsynonymous substitutions (dN), which are subject to selection. The ratio depends on variables such as effective population size and the divergence of the microbial populations compared (Kryazhimskiy and Plotkin, 2008). To determine the strength of selection on conserved and variable genes among the *Methanothermococcus* SAGs, we calculated ω for homologous ORF clusters containing sequences from two or more SAGs. ω values varied widely among the ORFs (Figure 5). In general, although ω values were somewhat higher than observed in some other microbial populations, e.g. (Hu and Blanchard, 2008; Luo et al., 2014; Martinez-Gutierrez and Aylward, 2019), The vast majority of ORFs had ω values less than 1, indicating purifying selection, which is consistent with previous work on signatures of selection in microbial genomes (Kryazhimskiy and Plotkin, 2008). However, several sets of gene clusters containing genes from SAGs in the C09/M21/K20 clade had elevated ω (closer to 1) (Figure 5). These gene clusters formed a group distinct from those that were shared between the C09/M21/K20 and E23/N22 clades (Figure 5). Very few SNPs were identified in this group of ORFs (highlighted in Figure 5), but the average ω for ORFs in this genomic region was close to 1 (Figure 5), indicating relaxed purifying selection or mild positive selection. Included in this group of ORFs were nitrogenases encoded on the genomic region that was unique to C09, M21 and K20.

**Figure 5.**
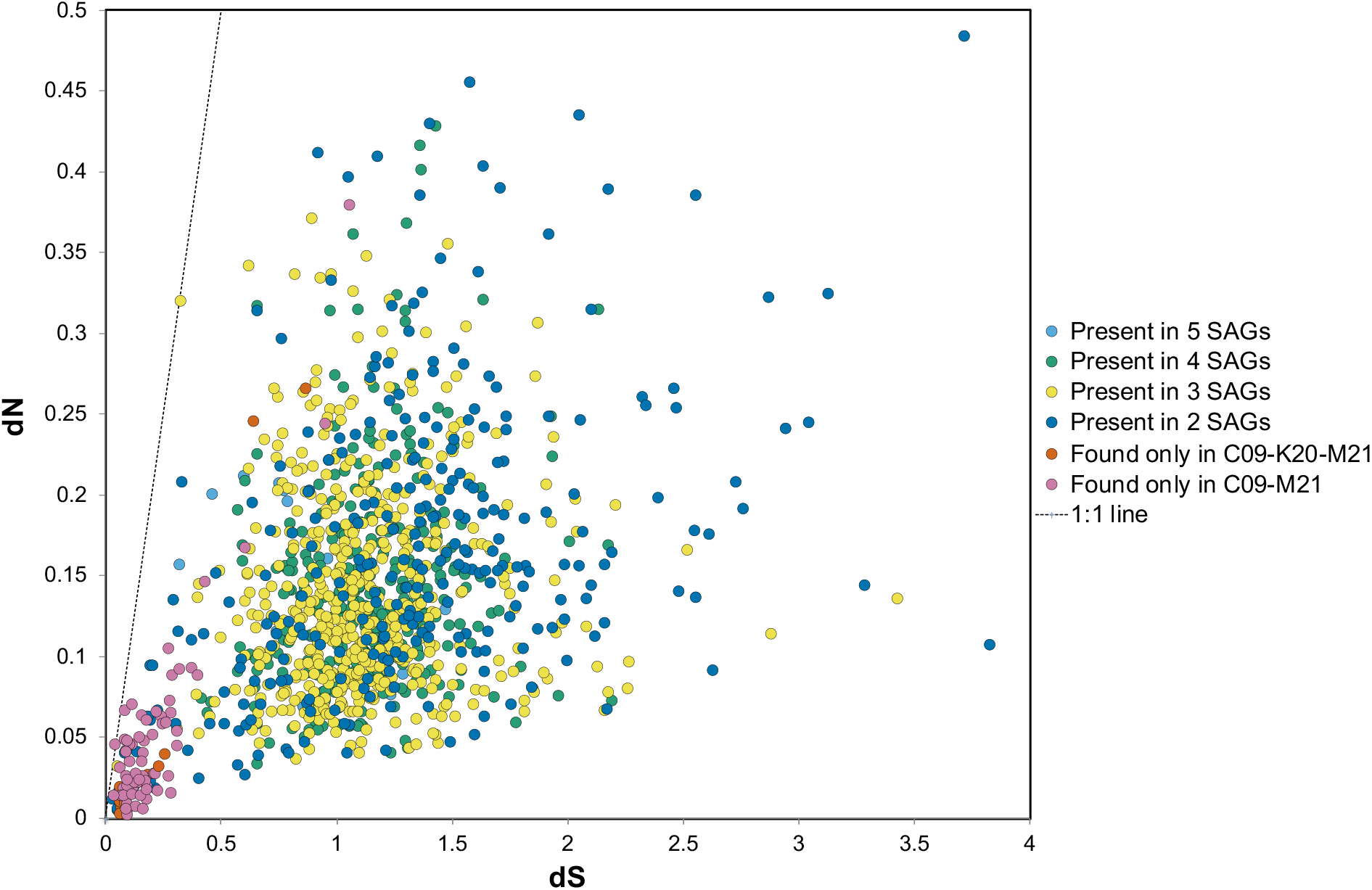
dN relative to dS for open reading frame (ORF) clusters. Light blue points correspond to ORF clusters found in all SAGs, green points are ORF clusters found in four SAGs, yellow in three, and dark blue in two. The dots in orange or pink represent ORFs shared between C09/M21 or C09/M21/K20, which includes the nitrogenase island of genes. Clusters of genes found in single genomes are excluded from this plot. The dotted line represents dN/dS = 1.

### Nucleotide variation within populations

Genomic variation within natural microbial populations can reveal whether specific populations have recently been subjected to selective sweeps or to clonal expansions (Bendall et al., 2016; Shapiro, 2016; Anderson et al., 2017). To characterize genomic variation within natural populations of *Methanothermococcus*, we measured single nucleotide variant (SNV) density and allele frequency (measured as n2n1) for each metagenome mapping to each SAG and compared this to coverage, which is a measure of relative abundance (Supplementary Figure 3). SNV density varied substantially within sample sites. Generally, SAGs with low average coverage had low SNV density (<10) and low n2n1 (<0.1) at those sample sites. In general, we consistently observed high SNV density associated with high coverage mappings (Figure S3). The relationship between mapping coverage and SNVs/kbp was positive, with R^2^ values above 0.80, indicating that increased mapping coverage also increased the number of SNVs observed (Supplementary Figure 4). However, the slope of the relationship varied between SAGs (Supplementary Figure 4), suggesting differential rates of SNV accumulation among the Methanothermococcus strains. SAGs with high phylogenetic relatedness showed similar trends of SNV density (Supplementary Figure 4). We did not observe cases in which SAGs had both high abundance and low SNV density, which would be interpreted as evidence of a clonal expansion or selective sweep.

## Discussion

Although methanogens are among the most common chemoautotrophic archaea observed in hydrothermal systems and constitute an important part of the microbial community, little is known about how they adapt and evolve in these globally distributed deep-sea ecosystems. Our results indicate that specific strains of *Methanothermococcus* have consistently higher genomic and transcript abundance in ultramafic hydrothermal vent settings (at Von Damm vent field) compared to mafic (at Piccard). Previous work has shown that methanogens in general, and *Methanothermococcus* in particular, were abundant in the Von Damm vent field (Reveillaud et al., 2016), and methanogens have been isolated from Von Damm (Sakai et al., 2019). The geochemistry of venting fluids at Von Damm is influenced by serpentinization, and thermodynamics calculations have suggested slightly higher catabolic energy availability for methanogenesis at Von Damm compared to Piccard (Reveillaud *et al*., 2016). Therefore, conditions at Von Damm are more favorable for elevated methanogen abundance and activity. In the present study, some *Methanothermococcus* lineages consistently displayed higher abundance relative to all others, indicating differences in fitness among the lineages represented by the SAGs, raising the question of what genomic features enabled some SAGs to have higher transcript abundance than others in these vent habitats. Although it is difficult to identify the specific genomic features that provide a selective advantage, notable differences among the strains studied here included genes related to nitrogen fixation, chemotaxis, CRISPR proteins and viruses. The genes and genomes also show distinct patterns of nucleotide variation across environments, suggesting that selection operates differently on these strains depending on the environmental context.

We identified several genomic features that may have contributed to the differences in fitness between strains. The E23 and N22 SAGs, which were more abundant than the other strains, were distinguished by the absence, rather the presence, of unique genes. In contrast, SAGs in the C09/K20/M21 clade shared a set of nitrogen-fixing genes in the *nif* family that are known to be part of the functional clade of cluster II nitrogenases. Diazotrophic growth is widespread among methanogens, including *Methanothermococcus* (Leigh, 2000; Nishizawa et al., 2014). Genomic islands encoding nitrogenase have previously been observed in the *Lebetimonas* (order *Nautiliales*) strains isolated from the subseafloor of hydrothermal vents (Meyer and Huber, 2014). In addition, nitrogen-fixing genes as well as demonstrated nitrogen fixing ability have been observed in methanogens from hydrothermal vents previously (Mehta et al., 2003; Mehta and Baross, 2006; Ver Eecke et al., 2013), including from a *Methanofervidicoccus* (order *Methanococcales)* strain isolated from the Mid-Cayman Rise (Wang et al., 2020). Although there is some ammonium enrichment relative to deep background seawater in Mid-Cayman Rise vent fluids, the maximum concentration is about 15-20 µmol, and nitrate is undetectable (Reeves et al., 2014; Charoenpong et al., 2015). Thus, fixed nitrogen sources are somewhat limited. Moreover, nitrogen-fixing genes are known to be part of the accessory genome in other microbial lineages in the vent environment (Meyer and Huber, 2014). The *nif* genes in these *Methanothermococcus* SAGs were transcribed and highly conserved. However, these genes were under relaxed purifying selection or mild positive selection, suggesting that natural selection allowed for some nucleotide diversification in these genes. Moreover, the nif genes do not appear to have provided a fitness advantage to these strains at the time of sampling, as SAGs encoding this genomic region had distinctly lower transcript abundance and lower overall abundance compared to the SAGs missing this genomic region. Thus, our results suggest that the possession of an accessory metabolic function (in this case, nitrogen fixation) does not necessarily confer an adaptive advantage to the strains possessing it with respect to population size, at least at the time of sampling.

Our results also indicate that *Methanothermococcus* strains in hydrothermal systems are actively infected by viruses. Previous research has shown that the genomes of thermophilic methanogens are particularly enriched with CRISPR loci (Anderson et al., 2011), and we identified several CRISPR loci in these SAGs. C09 and M21 encoded several distinct CRISPR-related proteins and lysogenic components of viruses, suggesting that these strains may be actively infected by viruses in the environment. CRISPR matches between the SAGs and the metagenomic contigs, transcripts of CRISPR and prophage-related genes, and the diversity and lack of conservation between spacers in the CRISPR loci all suggest that *Methanothermococcus* are susceptible to frequent viral infection in these vent systems. Previous work on the isolation of methanogen-targeting archaeal viruses from hydrothermal systems (Thiroux et al., 2020) and the high abundance of CRISPR loci in vent methanogens (Anderson et al., 2011) is consistent with this observation. However, we did not identify any CRISPR loci in the *Methanothermococcus* lineages with the highest abundance and gene transcription across vent sites (E23 and N22). It is possible that the CRISPR loci were not present due to incomplete sequencing and assembly, but the absence of any detectible CRISPR loci or CRISPR-related proteins on these two closely related strains suggests that, as with the nitrogenase cassette, possession of accessory functions such as CRISPR antiviral immune systems does not necessarily provide a selective advantage. Instead, both E23 and N22 encoded some restriction modification genes, membrane proteins, and glycosyltransferases that were absent in other strains. Both restriction enzymes and modifications to the cell wall and outer membrane pose alternative strategies by which these organisms may evade viral predation. Thus, the presence of antiviral mechanisms and variable CRISPR loci indicates that some of these strains may be actively infected by viruses in the environment, and that differential susceptibility to viral infection could contribute to differential fitness of the strains.

Finally, mapping of metagenomic reads to the SAGs allowed us to examine fine-scale variation at the nucleotide and amino acid levels, which revealed insights into selection pressure in the natural environment. We observed that closely related *Methanothermococcus* lineages accumulated variation at similar rates across different vent sites, indicating that there are consistent patterns of mutation accumulation that match *Methanothermococcus* phylogeny regardless of sample site. Thus, patterns of mutation accumulation or loss due to selection or drift appear to correlate closely to phylogeny. The density of SNVs was relatively high across all sample sites, suggesting that mutation rates were high or that clonal sweeps were not common in these methanogenic populations, in contrast to observations of microbial populations in previous studies (Bendall *et al*., 2016; Anderson *et al*., 2017). It appears that *Methanothermococcus* strains at these two hydrothermal vent sites are not susceptible to blooms or extinction events and are instead maintained in high abundances consistently over time, with specific strains maintaining higher abundance than others.

Together, our analysis of five *Methanothermococcus* SAGs alongside 15 metagenomes and 10 metatranscriptomes from diffuse flow fluids at the Mid-Cayman Rise reveals that the genomes of methanogens in vent environments exhibit a plasticity that can lead to differences in growth potential and gene expression across vent habitats. Some *Methanothermococcus* lineages exhibited higher gene and transcript abundance in some vent sites than in others. However, in contrast to observations of other strains in these vent fields (Anderson et al., 2017), we did not observe evidence for frequent clonal sweeps. Instead, the *Methanothermococcus* strains with the highest fitness appeared to have high rates of natural variation, indicating that they had been maintained in high abundance for long enough to accumulate variation. The most successful *Methanothermococcus* strains were notable for the absence of genes related to nitrogen fixation, prophage sequences and the CRISPR/Cas immune response, suggesting that the possession of accessory functions related to nitrogen fixation and viral immunity may not necessarily provide a selective advantage in these conditions. Continued work combining genomic and transcriptomic approaches will be key to elucidating the mechanisms by which methanogens co-evolve with their environment in hydrothermal vent systems.

## Acknowledgements

We would like to thank Victor Huerta and Jackson Raynor for assistance with bioinformatics analysis, and Chip Brier, Chris German, Jeff Seewald, Cindy Lee Van Dover, Jill McDermott, and Emily Reddington for assistance with sample collection, processing, and analysis. We would also like to thank the captains and crew of R/V *Atlantis* and R/V *Falkor* as well as ROVs *Jason* and *Nereus* for assistance with sample collection and preparation. We thank the SCGC staff for single cell genomics analyses. Support for M.H. was provided by Carleton College. RA was supported by a NASA Postdoctoral Fellowship with the NASA Astrobiology Institute. This work was supported by a grant from the Deep Carbon Observatory’s Census of Deep Life to RA and JAH, a NASA Exobiology grant number 80NSSC18K1076 to REA and JAH, a NASA Astrobiology Science and Technology for Exploring Planets (ASTEP) grant NNX-327 09AB75G to JAH, a grant from the Deep Carbon Observatory’s Deep Life Initiative to JAH and RS, the NSF Science and Technology Center for Dark Energy Biosphere Investigations (C-DEBI) and NSF grants OCE-1136488 and OIA-1826734 to RS. Data collected in this study in 2013 is based upon work supported by the Schmidt Ocean Institute during cruise FX008-2013 aboard R/V *Falkor*. Ship and vehicle time in 2012 were supported by the NSF-OCE grant OCE-1061863 to JSS. This is C-DEBI Contribution [number to be determined].

## Supplementary Figures

**Supplementary Figure 1.**
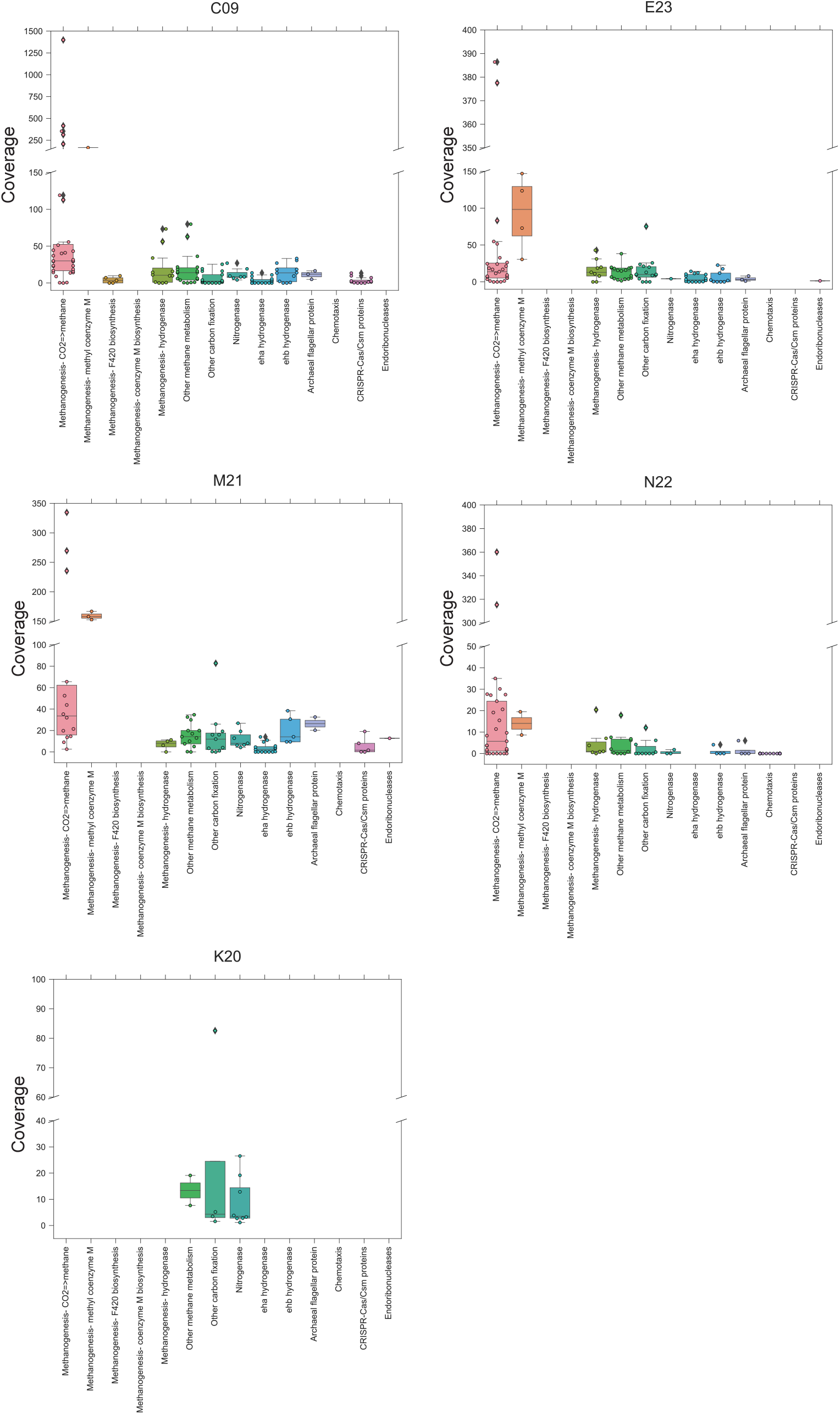
Box-and-whisker and swarm plots indicating the metatranscriptomic coverage of specific genes in *Methanothermococcus* single cell-amplified genomes in sample FS848, Ginger Castle, at Von Damm hydrothermal field. Coverage values are not normalized between SAGs, and thus relative transcript abundance of specific gene types should only be compared within individual SAGs. The boxes show the quartiles of the dataset, and the whiskers extend to show the rest of the distribution. Outliers are indicated by points outside of the whiskers. Individual dots indicate individual gene coverage values for genes within that category. Y axes are split to depict the full range of coverage values. A list of the KEGG numbers that were included within each category is listed in Supplementary Data File 1.

**Supplementary Figure 2.**
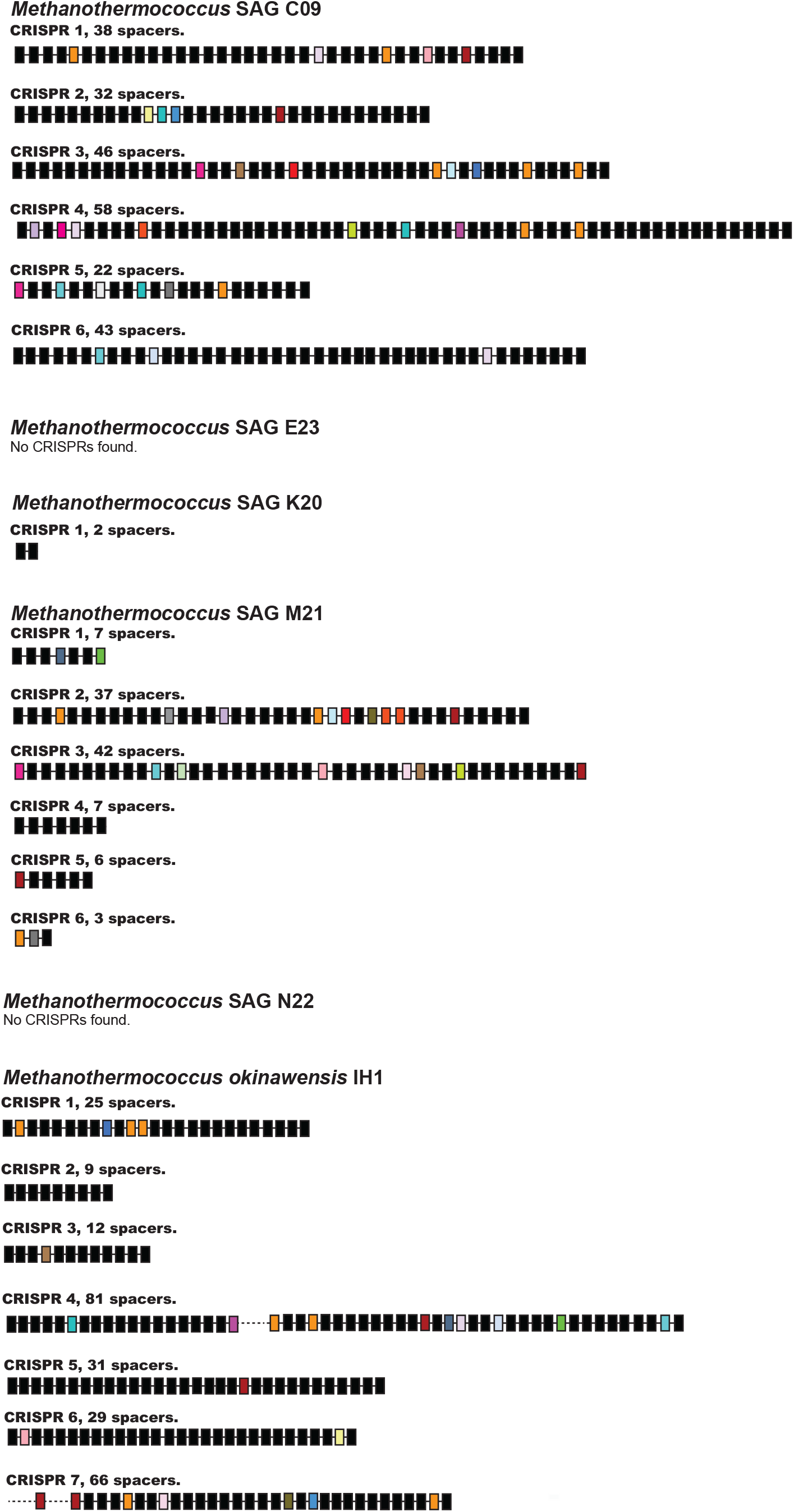
Summary of CRISPR loci in the 4 *Methanothermococcus* SAGs from Ginger Castle vent from Von Damm vent field on the Mid-Cayman Rise, sampled in 2012. Each of the CRISPR loci is represented by a line with a series of boxes, in which each box represents a spacer sequence. Matching spacers are shown in the same color; unique spacers are shown in black.

**Supplementary Figure 3.**
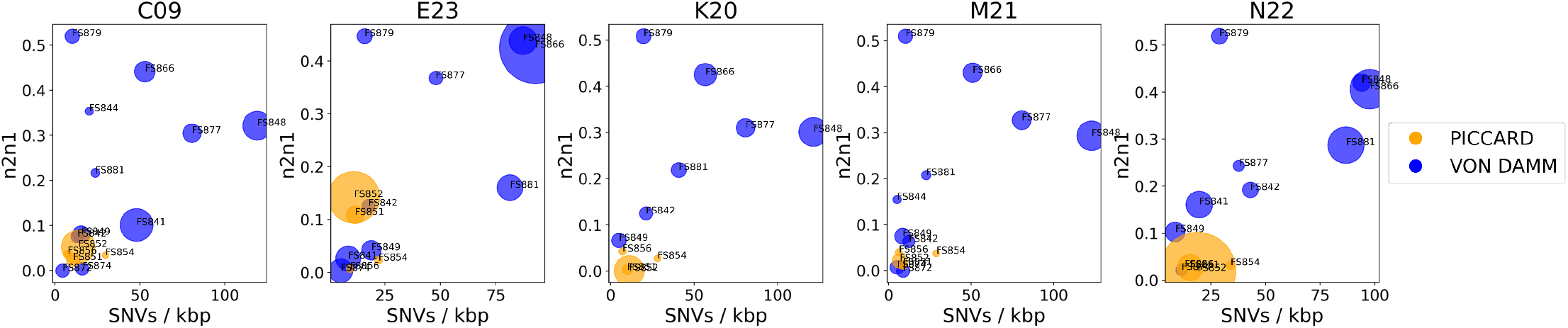
Plot of SNVs per kilobase pair versus n2n1 (majority allele frequency) for metagenome mappings to single cell amplified genomes from Piccard (blue-scale) and Von Damm (yellow-scale) vent fields. All y-axes reflect n2n1, or the ratio of the second most common allele to the most common allele. Bubble diameter corresponds to the average coverage of each SAG-metagenome mapping normalized by number of metagenomic reads. Bubbles are colored according to which vent field they came from; sample numbers are indicated.

**Supplementary Figure 4.**
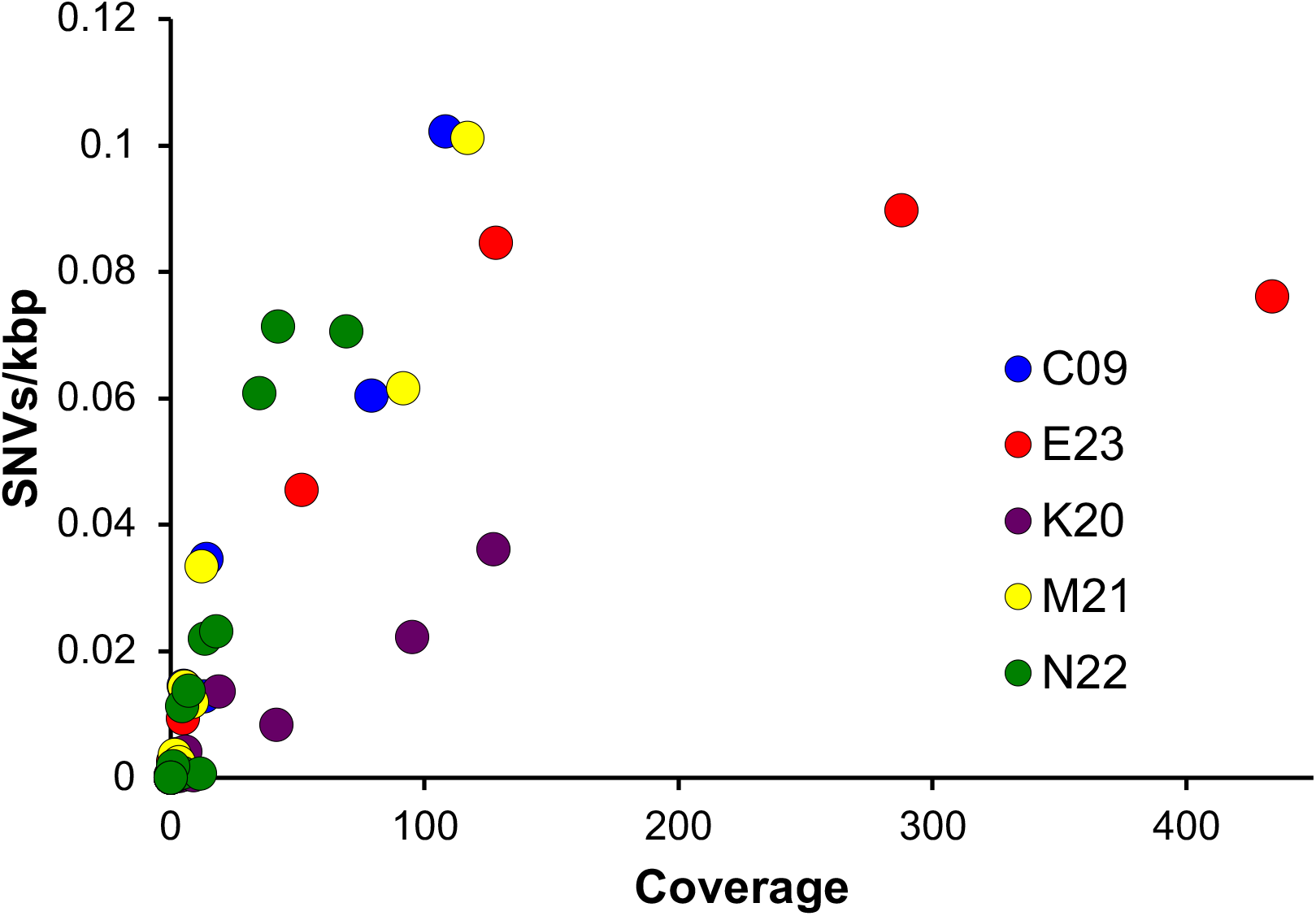
Scatterplots of average mapping coverage vs average number of SNVs per kilobase pair of DNA for metagenomic mappings, colored by SAG. Each point represents data from a different metagenome mapped to each SAG.

**Supplementary Table 1.** Data regarding quality filtering and assembly of metagenomic and metatranscriptomic reads. No MT, no metatranscriptome.

| Vent Field | Vent Name | Year | Sample Number | Latitude | Longitude | Temperature (°C) | Number of raw metagenomic reads (pairs) | Number of quality filtered metagenomic reads | ENA Sample Accession | ENA Secondary Accession | Number of raw metatranscriptomic reads (pairs) | Number of quality filtered metatranscriptomic reads, rRNA removed |
| --- | --- | --- | --- | --- | --- | --- | --- | --- | --- | --- | --- | --- |
| Von Damm | Old Man Tree | 2012 | FS841 | 18.375 | -81.8 | 114 | 61008797 | 53,972,022 | ERS1370013 | SAMEA4470834 | 14536191 | 3,427,865 |
| Von Damm | Ravelin #2 | 2012 | FS842 | 18.3752 | -81.8 | 86 | 69963233 | 64,064,672 | ERS1370014 | SAMEA4470835 | no MT | no MT |
| Von Damm | Shrimp Hole | 2012 | FS844 | 18.3747 | -81.8 | 50 | 111467935 | 104,733,336 | ERS1370015 | SAMEA4470836 | 44242399 | 29,768,330 |
| Von Damm | Ginger Castle | 2012 | FS848 | 18.3769 | -81.8 | 47 | 81658568 | 77,403,857 | ERS1370016 | SAMEA4470837 | 25427198 | 6,231,008 |
| Von Damm | Main Orifice | 2012 | FS849 | 18.3767 | -81.8 | 109 | 30239186 | 26,539,513 | ERS1370017 | SAMEA4470838 | no MT | no MT |
| Piccard | Hot Chimlet, BVM | 2012 | FS851 | 18.5472 | -81.72 | 106 | 88561544 | 78,203,317 | ERS1370018 | SAMEA4470839 | 25550688 | 5,385,293 |
| Piccard | Shrimp Canyon, BVM | 2012 | FS852 | 18.5471 | -81.72 | 44 | 42204178 | 38,058,339 | ERS1370019 | SAMEA4470840 | 26316504 | 10,571,338 |
| Piccard | X-19 at BV #4, BVM | 2012 | FS854 | 18.5466 | -81.72 | 18 | 148334278 | 141,073,647 | ERS1370020 | SAMEA4470841 | 12643619 | 1,126,747 |
| Piccard | Shrimp Gulley #2, BSM | 2012 | FS856 | 18.5466 | -81.72 | 108 | 109104481 | 104,273,127 | ERS1370021 | SAMEA4470842 | 11471361 | 3,822,920 |
| Von Damm | near Main Orifice | 2013 | FS866 | 18.3767 | -81.8 | 130 | 27575200 | 25,829,542 | ERS1370022 | SAMEA4470843 | no MT | no MT |
| Von Damm | Shrimp Hole | 2013 | FS872 | 18.3747 | -81.8 | 30 | 45935925 | 40,627,197 | ERS1370023 | SAMEA4470844 | 22530122 | 19,601,515 |
| Von Damm | Twin Peaks | 2013 | FS874 |  |  | 140 | 72719556 | 67,240,297 | ERS1370024 | SAMEA4470845 | no MT | no MT |
| Von Damm | Shrimp Buttery | 2013 | FS877 |  |  | 131 | 167341615 | 158,036,033 | ERS1370025 | SAMEA4470846 | no MT | no MT |
| Von Damm | Hot Cracks #2 | 2013 | FS879 | 18.375 | -81.8 | 29 | 27319409 | 25,899,380 | ERS1370026 | SAMEA4470847 | 21345009 | 8,229,809 |
| Von Damm | Old Man Tree | 2013 | FS881 | 18.375 | -81.8 | 114 | 191547095 | 178,390,689 | ERS1370027 | SAMEA4470848 | 14231356 | 8,856,348 |

**Supplementary Table 2.** Percent reads mapping for all SAG-to-SAG mappings. SAG contigs used as reference are listed on the left, SAG reads mapped to the reference are listed along the top of the table. Numbers are listed as percent of merged reads mapped using bowtie2.

| SAG reference | SAG reads |  |  |  |  |  |
| --- | --- | --- | --- | --- | --- | --- |
|  |  | C09 | E23 | K20 | M21 | N22 |
|  | C09 | 95.82 | 0 | 33.83 | 64.14 | 0.1 |
|  | E23 | 0 | 93.43 | 0 | 0 | 1.03 |
|  | K20 | 4.6 | 0 | 97.7 | 2.11 | 0 |
|  | M21 | 42.44 | 0 | 18.3 | 96.53 | 0.05 |
|  | N22 | 1.09 | 2.15 | 1.23 | 0.08 | 65 |

**Supplementary Table 3.** Selected gene counts for genes present and/or absent in each SAG from the Mid-Cayman Rise.

| Gene name | Category | C09 | E23 | K20 | M21 | N22 |
| --- | --- | --- | --- | --- | --- | --- |
| iron transport protein (FeoB_CN) | **Fe** | 2 | 3 | 0 | 0 | 2 |
| iron transport protein (FeoA) | **Fe** | 3 | 4 | 0 | 1 | 2 |
| ech hydrogenase (echABCE) | **H<sub>2</sub>** | 0 | 0 | 0 | 0 | 0 |
| energy-converting hydrogenase A (ehaBCEGNOP) | **H<sub>2</sub>** | 28 | 24 | 0 | 20 | 10 |
| energy-converting hydrogenase B (ehbABFIJLNO) | **H<sub>2</sub>** | 28 | 24 | 0 | 20 | 10 |
| coenzyme F420 hydrogenase (frhAGDG) | **H<sub>2</sub>** | 2 | 2 | 0 | 1 | 2 |
| hydrogenase (hyaABC, hybO, hybC) | **H<sub>2</sub>** | 0 | 0 | 0 | 0 | 0 |
| [NiFe] hydrogenase (hydA2A3B2B3) | **H<sub>2</sub>** | 0 | 0 | 0 | 0 | 0 |
| methane/ammonia monooxygenase (pmoA-amoA) | **CH<sub>4</sub>** | 0 | 0 | 0 | 0 | 0 |
| methyl-coenzyme M reductase (mcrABCDG) | **CH<sub>4</sub>** | 3 | 3 | 0 | 3 | 4 |
| cytochrome c oxidase (coxABCD,AC,ctaF) | **O<sub>2</sub>** | 0 | 0 | 0 | 0 | 0 |
| cytochrome c oxidase cbb3-type (ccoNOPQ_NO) | **O<sub>2</sub>** | 0 | 0 | 0 | 0 | 0 |
| carbon-monoxide dehydrogenase (cooCFS, acsA) | **O<sub>2</sub>** | 2 | 2 | 0 | 0 | 0 |
| fumarate reductase (frdAB) subunitAB | **C** | 2 | 2 | 1 | 2 | 2 |
| Nitrogenase | **N** | 1 | 0 | 1 | 1 | 0 |
| Phosphoadenosine phosphosulfate reductase/PAPS reductase | **S** | 1 | 1 | 0 | 1 | 0 |
| Serine O-acetyltransferase (cysE) | **S** | 1 | 1 | 1 | 0 | 1 |
| Chemotaxis proteins (M. okinawensis 1415, 1417, 1418; chemotaxis proteins CheW, CheY and CheA; methyl-accepting chemotaxis signaling domain-containing protein, methyl-accepting chemotaxis sensory transducer) | **Chemo-taxis** | 0 | 0 | 0 | 0 | 7 |
| Cas proteins (cas1-6) | **CRISPR** | 6 | 0 | 0 | 0 | 0 |
| Csm proteins (csm1-5) | **CRISPR** | 5 | 0 | 0 | 5 | 0 |

**Supplementary Data File 1:** An Excel spreadsheet that lists all ORF clusters as well as a list of the KO categories used to make the box-and-whisker plot.

